# A dynamic role for dopamine receptors in the control of mammalian spinal networks

**DOI:** 10.1101/715326

**Authors:** Simon A. Sharples, Nicole E. Burma, Joanna Borowska-Fielding, Charlie H.T. Kwok, Shane E.A. Eaton, Glen B. Baker, Celine Jean-Xavier, Ying Zhang, Tuan Trang, Patrick J. Whelan

## Abstract

Dopamine is well known to regulate movement through the differential control of direct and indirect pathways in the striatum that express D_1_ and D_2_ receptors respectively. The spinal cord also expresses all dopamine receptors however; how the specific receptors regulate spinal network output in mammals is poorly understood. We explore the receptor-specific mechanisms that underlie dopaminergic control of spinal network output of neonatal mice during changes in spinal network excitability. During spontaneous activity, which is a characteristic of developing spinal networks operating in a low excitability state, we found that dopamine is primarily inhibitory. We uncover an excitatory D_1_-mediated effect of dopamine on motoneurons and network output that also involves co-activation with D_2_ receptors. Critically, these excitatory actions require higher concentrations of dopamine; however, analysis of dopamine concentrations of neonates indicates that endogenous levels of spinal dopamine are low. Because endogenous levels of spinal dopamine are low, this excitatory dopaminergic pathway is likely physiologically-silent at this stage in development. In contrast, the inhibitory effect of dopamine, at low physiological concentrations is mediated by parallel activation of D_2_, D_3_, D_4_ and α_2_ receptors which is reproduced when endogenous dopamine levels are increased by blocking dopamine reuptake and metabolism. We provide evidence in support of dedicated spinal network components that are controlled by excitatory D_1_ and inhibitory D_2_ receptors that is reminiscent of the classic dopaminergic indirect and direct pathway within the striatum. These results indicate that network state is an important factor that dictates receptor-specific and therefore dose-dependent control of neuromodulators on spinal network output and advances our understanding of how neuromodulators regulate neural networks under dynamically changing excitability.

**Significance statement:** Monoaminergic neuromodulation of neural networks is dependent not only on target receptors but also on network state. We studied the concentration-dependent control of spinal networks of the neonatal mouse, in vitro, during a low excitability state characterized by spontaneous network activity. Spontaneous activity is an essential element for the development of networks. Under these conditions, we defined converging receptor and cellular mechanisms that contribute to the diverse, concentration-dependent control of spinal motor networks by dopamine, in vitro. These experiments advance understanding of how monoamines modulate neuronal networks under dynamically changing excitability conditions and provide evidence of dedicated D_1_ and D_2_ regulated network components in the spinal cord that are consistent with those reported in the striatum.

## Introduction

Neuromodulators are critical for central nervous system function and diversify circuit outputs by altering synaptic and intrinsic properties [1–3]. Dopamine is a monoamine neuromodulator that is well known for action selection in vertebrates through the control of direct and indirect circuits of the basal ganglia that express excitatory D_1_ and inhibitory D_2_ receptors respectively (for review, see [4] and [5–7] for examples). Dopamine is also important for the regulation of spinal motor networks that control rhythmic movements including but not limited to locomotion (for review, see [8]). In larval zebrafish, phasic and tonic firing patterns of descending neurons that provide the primary source of spinal dopamine correlates with locomotor episodes and quiescence, respectively [9]. These different firing patterns can impact the cellular release of dopamine and as a consequence, the receptor subtypes it activates [10]. For example, high levels of dopamine released during phasic cell firing activate lower affinity excitatory D_1_ receptors and promote locomotor activity, and lower levels of dopamine released during tonic activity activate higher affinity inhibitory D_2_ receptors and suppress motor output [11]. Similarly, dopamine has dose-dependent effects on locomotor circuits in precocial species with functional swim networks [12].

We have recently demonstrated that neuromodulation of developing mammalian spinal circuits is state-dependent [13] which is consistent with work in invertebrates [14,15]. This is important in developing spinal motor networks which produce a wide repertoire of patterned outputs at birth, including locomotor activity, as a consequence of dynamically fluctuating network excitability. That being said, neonatal rodents rarely produce coordinated bouts of locomotion and in vitro preparations of isolated spinal cord require pharmacological or electrical stimulation to drive the network into a high excitability state to produce fictive walking patterns. Instead, most of the movements observed in neonatal mice are ataxic. (See Supplementary Video 1). These movements correlate with spontaneous network activity, which can be observed in vitro [16–19]. Nevertheless, the vast majority of what we know about neuromodulation of developing mammalian spinal networks has been derived from studies on fictive locomotor activities with dopamine being predominantly excitatory when spinal networks are operating in this state [20–31].

Based on our previous work, we hypothesized that receptor actions, and therefore concentration-dependent control of network output by dopamine, is linked to the underlying network excitability state [13]. As a result, more complex receptor-dependent actions may have been masked during high excitability states of fictive locomotion. We therefore examined how dopamine modulates spinal output at the network and cellular level of neonatal mouse spinal cords in vitro during a low excitability state characterized by spontaneous activity [16,17]. We found that the receptor-specific effects of dopamine are fundamentally different during a low excitability state compared to what has been previously reported during high excitability states. Specifically, during a low excitability state, dopamine has primarily inhibitory effects on network output acting in parallel through activation of D_2_, D_3_, D_4_, and α_2_ receptors. We uncover an excitatory effect of dopamine that is likely physiologically silent. This is because endogenous dopamine levels in the spinal cord are low at this age and the excitatory effects, involving coactivation of both D_1_ and D_2_ receptors, require higher concentrations of dopamine. We also found that excitatory and inhibitory dopaminergic pathways act through the control of dedicated network components. Specifically, the D_1_ pathway acts through excitation of motoneurons, likely recruiting recurrent excitatory circuits [32], with the D_2_ pathway acting through hyperpolarization of multiple ventral interneuron subclasses. Portions these data were presented in abstract form [33,34].

## Results

### Dopaminergic modulation of spinal motor networks is dose-dependent and bidirectional

We focused on the modulation of perinatal spontaneous activity patterns to investigate how dopamine modulates spinal network output during a physiologically relevant low excitability state. Using the same experimental set-up as Sharples et al. [23], we recorded spontaneous motor activity with extracellular suction electrodes from single ventral roots of the second or fifth lumbar (L2/L5) segments simultaneously from two or four spinal cord preparations sharing the same chamber. Each preparation in this series of experiments was naïve to dopamine exposure and received only a single dose of its respective concentration. This configuration ensured consistent experimental conditions across several preparations exposed to different concentrations of dopamine. Low concentrations of dopamine (1–30 µM) consistently suppressed spontaneous motor activity, whereas higher concentrations (100–300 µM) excited spontaneous motor activity, evoking episodic and continuous rhythmic patterns (Fig. 1; n = 42, F_(5,36)_ = 10.5. p < 0.001).

**Figure 1:**
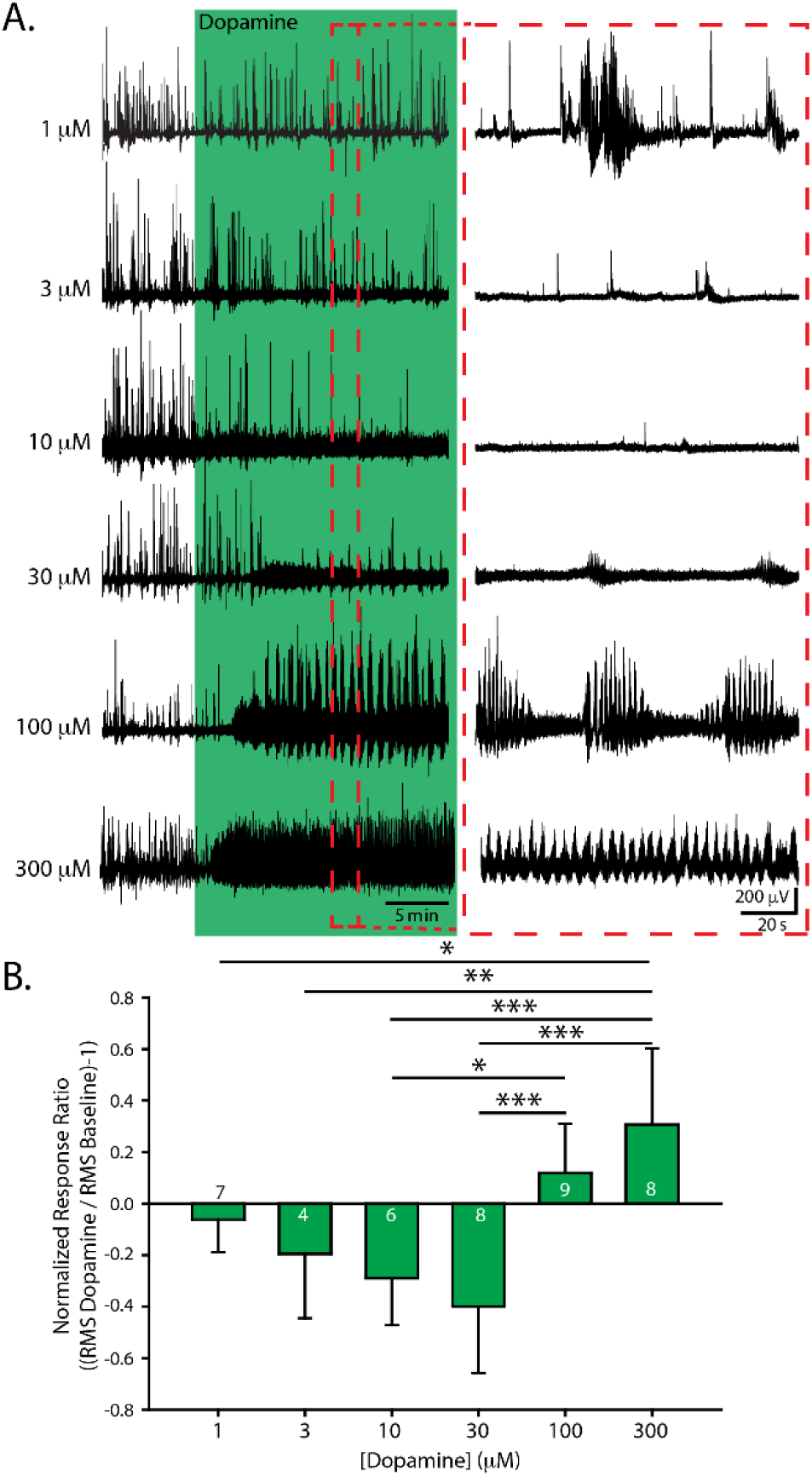
Dopamine evokes bidirectional dose-dependent modulation of lumbar network activity. A. Extracellular neurograms recorded from naïve single lumbar ventral root preparations after applying dopamine in various concentrations. Each row indicates a separate trial. Each recording began with spontaneous motor activity; the green section shows the effect of dopamine for each dose. The red-dashed box highlights expanded sections of data to the right of each neurogram. B. Mean normalized response ratios calculated from the root mean square of the raw neurogram during a 5-minute window, 20 minutes after dopamine application, compared to baseline activity. The negative response ratios represent inhibition and positive values represent excitation. The number in each green bar indicates the number of preparations in the average for each concentration. Error bars indicate standard deviation. Asterisks denote the significance level of Tukey post hoc … … tests between ratio concentrations (*p < 0.05, **p < 0.01, ***p < 0.001).

### Parallel actions of D_2_, D_3_, D_4_, dopamine and alpha-2 adrenergic receptors mediate dopaminergic inhibition of spontaneous activity

To delineate receptor contributions to dopamine’s bidirectional effects on endogenous spontaneous activity in isolated spinal cord preparations, we used antagonists selective for the family of dopamine receptors. At low concentrations of dopamine, we observed a negative response ratio (Fig. 2A, Di), which was due to a reduction in the number (Fig. 2Dii), but not the amplitude, of spontaneous episodes (Fig. 2Diii). In contrast to our hypotheses, the inhibitory effect of dopamine at low concentrations (10 µM) was not altered by antagonists targeting D_2_ (L-741,626; n = 3, response ratio = −0.57 ± 0.26), D_2_/D_3_ (sulpiride; 20 µM; n = 10, response ratio = −0.36 ± 0.2), D_3_ (SB-27701-A, 5 µM; n = 3; response ratio = −0.36 ± 0.3), D_4_ (L-745,870; n = 3, response ratio = −0.32 ± 0.1) or D_1_/D_5_ receptors (SCH 23390; n = 4, response ratio = −0.43 ± 0.2). However, when all D_2_-like receptors were blocked with a cocktail of sulpiride (D_2_/D_3_, 20 µM) and L-745,870 (D_4_, 5 µM), response ratios indicated that the inhibitory effect of 10 µM dopamine was attenuated, compared with dopamine alone (Fig. 2C; n = 6; one-way analysis of variance (ANOVA), F_(4,32)_ = 7.3, p < 0.001; Tukey post hoc p = 0.09), to a level where it was not significantly different from a time-matched vehicle control (Fig. 2C, Di.; n = 6, F_(4,32)_ = 7.3, p < 0.001; Tukey post hoc, p = 0.98). Burst analysis revealed no difference in the number or amplitude of episodes when antagonists were present, compared with baseline (Fig. 2Dii, Diii; two-way ANOVAs for number, F_(2,28)_ = 9.5, p < 0.0001; two-way ANOVA for amplitude: F_(4,28)_ = 3.4, p = 0.023). Together these data suggest that the inhibitory actions of dopamine depend on parallel action of D_2_, D_3_ and D_4_ receptors.

**Figure 2:**
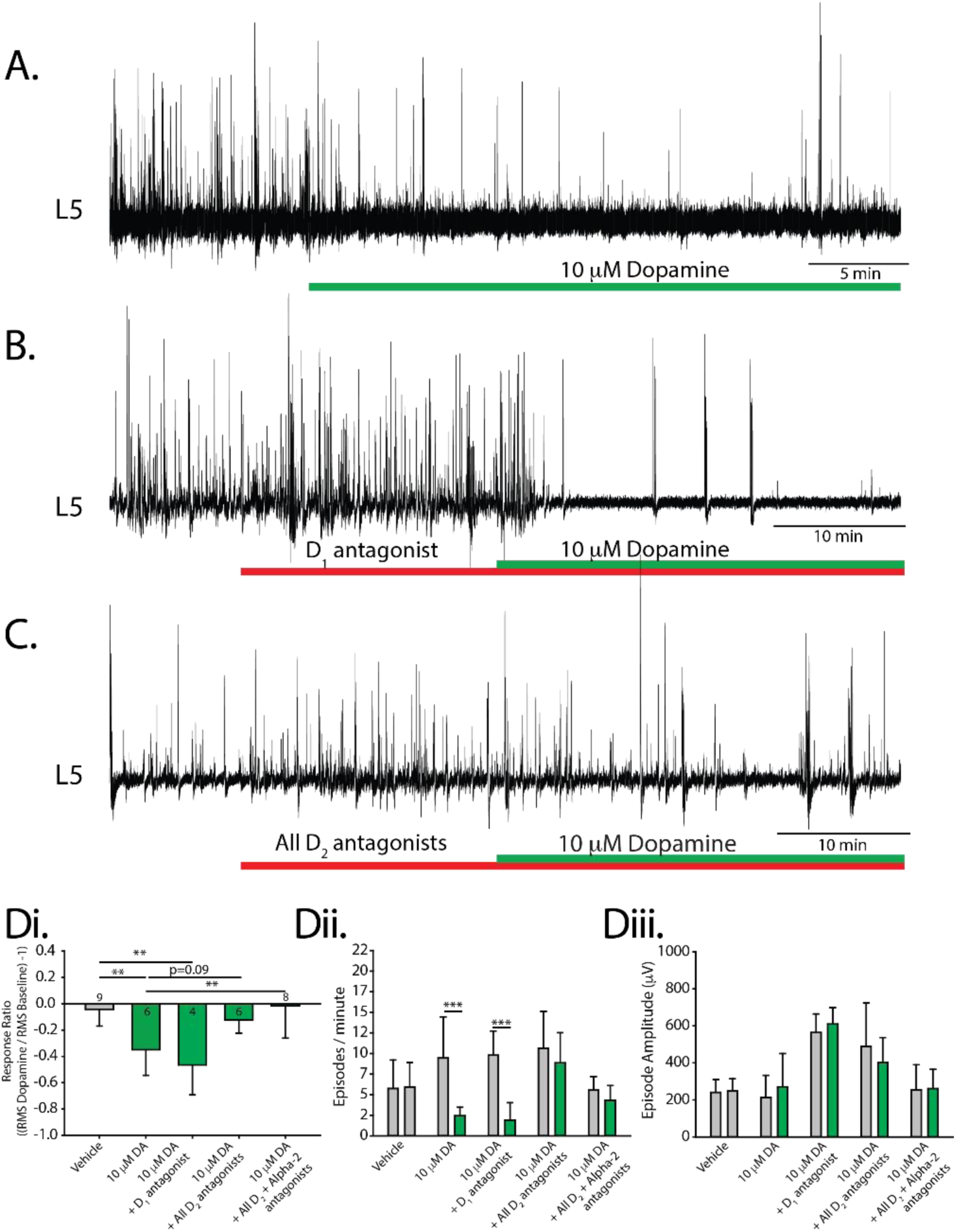
Receptor mechanisms for dopaminergic inhibition of spontaneous network activity. A–C. Single ventral root (L5) extracellular neurograms of spontaneous activity in the presence of dopamine (A), a D_1_ antagonist and dopamine (B), and with a cocktail of D_2_ antagonists and dopamine (C). Spinal cords were perfused with receptor-preferring antagonists (red bars) 20 minutes prior to the application of low concentrations of dopamine (10 µM). D. Response ratio represents the root mean square of raw neurograms during a 5-minute window, 20 minutes after dopamine application, compared with prior to dopamine application. Negative response ratios indicate inhibition and positive values indicate excitation. The number in each bar indicates the number of preparations included in that condition’s mean. Bars indicate the mean (± SD), number of episodes per minute (Dii), and amplitude (Diii) of spontaneous episodes recorded within the 5-minute epochs during which response ratios were calculated. Asterisks indicate the significance level of post hoc comparisons (*p < 0.05, **p < 0.01, ***p < 0.001).

We considered the possibility that the inhibitory influence of low dopamine concentrations was partly due to the activation of non-dopamine receptors. Previous work conducted in our laboratory showed that dopamine inhibits cauda equina-evoked locomotion, partially via α_2_-adrenergic receptors [30]. The remaining minor inhibitory effect of dopamine at 10 µM was blocked by antagonizing the α_2_-adrenergic receptors with yohimbine (4 µM) in the presence of D_2_/D_3_ and D_4_ receptor antagonists. We observed significantly different response ratios, compared with 10 µM dopamine alone (Fig. 2Di; n = 6, F_(4,32)_ = 7.3, p < 0.001; Tukey post hoc, p = 0.006). Low dopamine concentrations did not change the number or amplitude of episodes in the presence of these antagonists, compared with baseline (Fig. 2Dii, Diii) suggesting that non-dopamine receptors also contribute to dopamine’s inhibitory effects.

### D_1_ - D_2_ receptor coactivation contributes to dopaminergic excitation of spinal network activity

Dopamine binding to D_1_ and D_2_ heteromers can lead to depolarization via an increase in intracellular calcium levels, mediated by the enzyme phospholipase C (PLC) [35–38]. Thus, we first tested the role of the lower affinity D_1_ receptor system and then examined whether the D_2_ receptor system has a cooperative role in the control of spontaneous activity. With the addition of dopamine at high concentrations (i.e., 50 & 100 µM), spontaneous activity patterns became rhythmic, often producing a slow rhythm with episodes of high frequency rhythmic activity. Moreover, the presence of the D_1_-like receptor antagonist, SCH-23390, reduced this effect (Fig. 3; 100 µM dopamine with 10 µM SCH-23390; n = 5; response ratio, F_(5,35)_ = 11.4, p < 0.001); fast rhythm power, F_(4,27)_ = 12.6, p < 0.001; slow rhythm power, H_(4)_ = 12.8, p = 0.013; Fig. 3 B & C, 50 µM dopamine with10 µM SCH-23390; response ratio, F_(5,35)_ = 11.4, p < 0.001; fast rhythm power, F_(4,27)_ = 12.6, p < 0.001; slow rhythm power, H_(4)_ = 12.8, p = 0.013, Dunn’s post hoc: p = 0.03; one way ANOVA used in tests). These results suggest that the excitatory effects of dopamine are primarily mediated by the D_1_-like receptor family.

**Figure 3:**
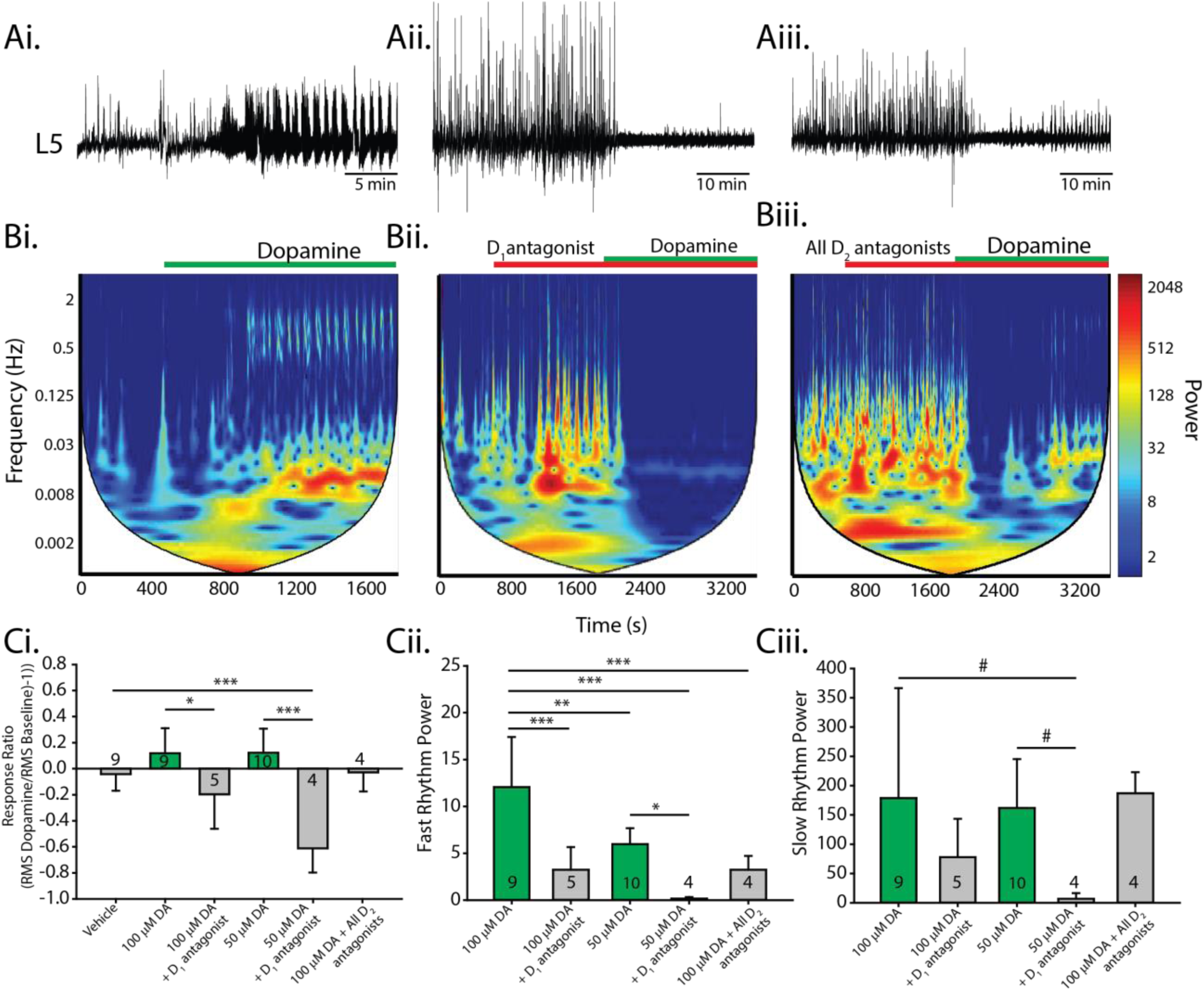
Receptor mechanisms for dopaminergic excitation of spinal network activity. High concentrations of dopamine excite spinal networks and produce episodic and continuous rhythmic patterns of activity. A. Single ventral root extracellular neurograms of spinal network activity from each condition: dopamine applied alone (Ai), in the presence of a D_1_ antagonist (Aii), and with a cocktail of D_2_ antagonists (Aiii). Horizontal bars indicate the timing of dopamine application (green bars) and the application of receptor-preferring antagonists (red bars). B. Spectrograms show autowavelet frequency power across time, evoked at high concentrations of dopamine (50-100 µM). The colour bar indicates power magnitude, from high (warm) to low (cool colours). We selected regions of interest around regions that coincided with fast and slow rhythm frequency rhythms. Spinal cords were perfused with antagonists for 20 minutes prior to the application of dopamine. Ci. Mean response ratios (± SD), as in Fig. 2, for each condition. Spectral analysis of fast rhythm (Cii) and slow rhythm (Ciii) power following drug application for each experimental condition. Histograms present mean values ± SD, with asterisks denoting significance level of post hoc tests (*p < 0.05, **p < 0.01, ***p < 0.001) following one-way ANOVA (fast rhythm; Cii) or nonparametric one-way ANOVA (slow rhythm; Ciii).

If D_2_ receptors form heteromers with D_1_ receptors, we would expect that antagonists would inhibit, rather than excite responses. We tested this idea by administering D_2_-like antagonists (sulpiride + L745,870) and found that the power of the fast rhythm elicited by dopamine at 100 µM was reduced (Fig. 3 Aiii, Biii, Cii; n = 4; F_(4,27)_ = 12.6, p < 0.001) to the same extent as the D_1_-antagonist, with no effect on the power of the slow rhythm (Fig. 3Ciii; H_(4)_ = 12.8, p = 0.013; Dunn’s post hoc, p= 1.0). This suggests that D_2_ receptors may contribute to the excitatory effects of dopamine.

To explore the interaction and co-activation profile of D_1_ and D_2_ receptors in isolated neonatal spinal cords, we performed co-immunoprecipitation for D_1_ and D_2_ receptors and used agonists to activate both receptor subtypes. After immunoprecipitating D_2_ receptors from neonatal spinal cord lysates, we used an antibody to probe for D_1_ receptors. We detected D_1_ receptor protein within the D_2_ receptor immunoprecipitates (Fig. 4Ciii) and these bands were blocked when pre-incubated with an antigen-blocking peptide. This result indicates that D_1_ and D_2_ receptors form a protein complex in neonatal mouse spinal cords.

**Figure 4:**
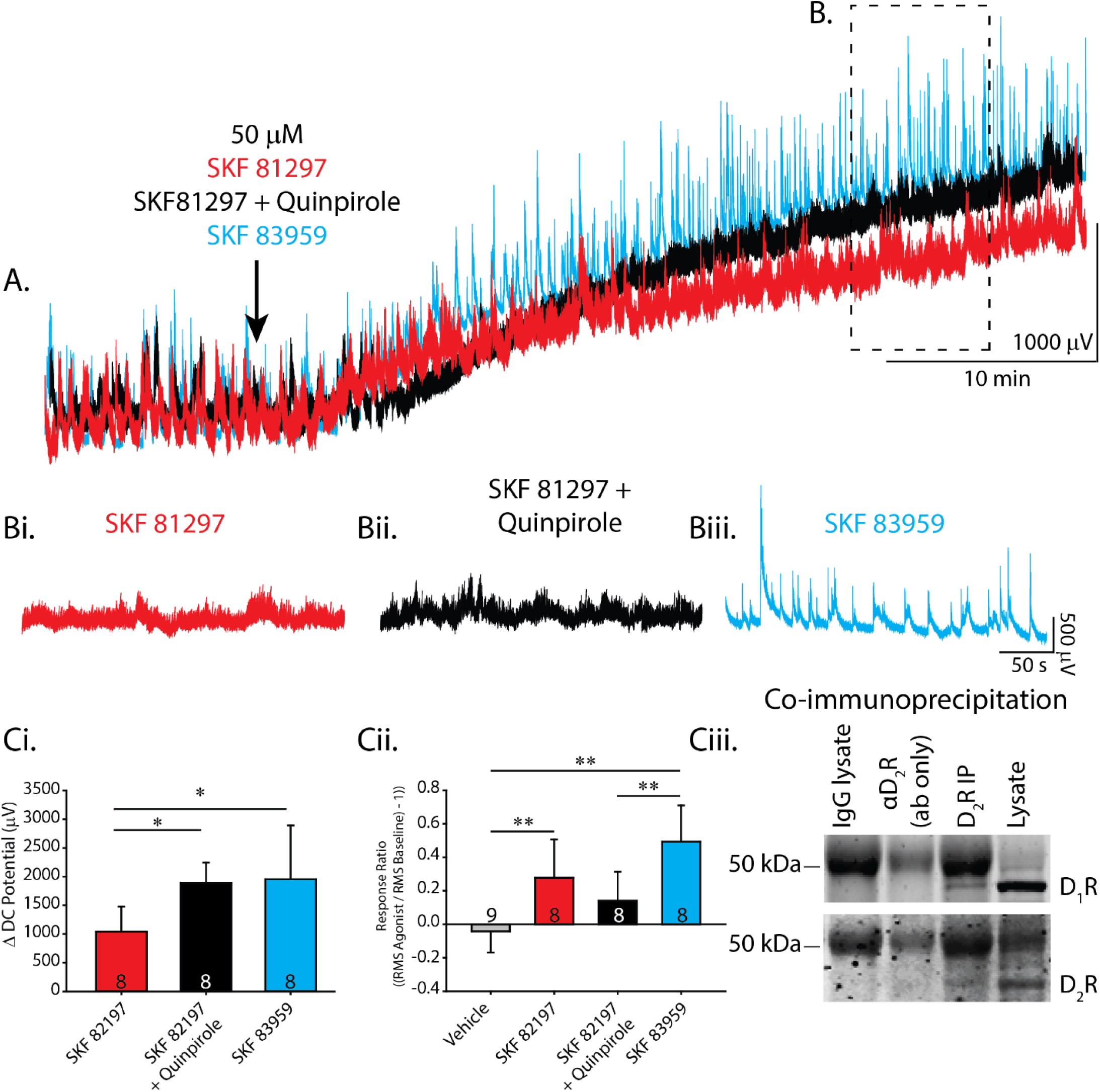
D_1_ D_2_ receptor co-activation contributes to the excitatory effect of dopamine. A. Single ventral root extracellular DC neurograms of spontaneous activity from conditions where a D_1_ agonist (SKF 81297) was applied alone (red), co-applied with a D_2_ agonist (SKF 81297 + Quinpirole; black), or D_1_ and D_2_ receptors were coactivated by a D_1_/D_2_ co-agonist (SKF 83959; blue). Bi-iii. For each agonist, (B) expands the region of spontaneous activity 20–25 minutes after agonist application, as outlined by the dashed line in (A). C. Ci depicts depolarization of DC potentials from DC neurograms, averaged across the number of preparations denoted by numbers in each condition’s bar. Cii shows the response ratio for each condition (as in previous figures), which represents changes in the amount of spontaneous activity. Ciii. Co-immunoprecipitation of D_1_ with D_2_ receptors suggests that they may interact in the neonatal spinal cord. Histograms present mean values ± SD, with asterisks denoting significance level of post hoc comparisons (*p < 0.05, **p < 0.01, ***p < 0.001) following one-way ANOVA.

In support of this idea, co-application of the D_1_ agonist SKF 81297 (50 µM) and the D_2_ agonist quinpirole (50 µM) elicited a more robust depolarization of the ventral root DC potential, compared with 50 µM of the D_1_ agonist alone (Fig. 4A, Ci; D_1_, n = 8; D_1_/D_2_, n = 8; one-way ANOVA F_(2,21)_ = 5.2, p = 0.01; Tukey post hoc: p = 0.02). We observed no difference in the amount of spontaneous network activity evoked with co-application of a D_2_ agonist, compared with application of the D_1_ agonist alone, as indicated by the response ratio (Fig. 4B, Cii; one-way ANOVA, F_(3,29)_ = 12.0, p < 0.001; Tukey post hoc, p = 0.5). In contrast, lower concentrations of the same agonists (10 µM) produced no effects (n = 8 for each condition; DC potential, t_(6)_ = 0.73, p = 0.24; response ratio, t_(6)_ = 0.9, p = 0.19). Thus, consistent with previous reports for striatal neurons [36], we found a dose-dependent effect of dopamine agonists wherein co-applying high doses, but not low doses, of D_1_ and D_2_ receptor agonists, produced more robust depolarization than a D_1_ agonist alone.

In addition to co-applying separate D_1_ and D_2_ agonists, we tested co-activating D_1_ and D_2_-like receptors with the D_1_/D_2_ co-agonist SKF 83959 (50 µM) [38,39]. As predicted, the co-agonist elicited a more robust depolarization of the ventral root DC potential, compared with the D_1_ agonist, when applied alone (Fig. 4A, Ci; D1, n = 8; D_1_/D_2_, n = 8; one-way ANOVA, F_(2,21)_ = 5.2, p = 0.01; Tukey post hoc, p = 0.03). Interestingly, the co-agonist also robustly facilitated superimposed spontaneous activity, as indicated by a larger response ratio than co-application of the D_1_ and D_2_ agonists produced (Fig. 4B, Cii; one-way ANOVA, F_(3,29)_ = 12.0, p < 0.001; Tukey post hoc, p = 0.004). These data suggest that under certain conditions, D_2_ receptors that are typically inhibitory can play an excitatory role via D_1_/D_2_ co-activation and contribute to lumbar motor network excitation in the neonatal mouse spinal cord.

### Low levels of endogenous spinal dopamine inhibit spontaneous activity

We next examined how the endogenous dopamine system regulates perinatal spinal network function. Endogenous levels of dopamine were increased by blocking the dopamine transporter (DAT) with an antagonist GBR 12909 (10 µM). Blocking DAT (n = 8 preparations) produced a modest but significant reduction in spontaneous activity compared to time-matched vehicle controls, reflected by a reduced response ratio (Fig. 5A, Bi; one-way ANOVA, F_(2,19)_ = 18.0, p < 0.001) and a reduced number of spontaneous episodes (Fig. 5Bii; n = 8, two-way ANOVA, F_(2,20)_ = 11.8, p = 0.0004), with no change in amplitude (Fig. 5Biii; n = 6, two-way ANOVA, F_(2,20)_ = 1.3, p = 0.3). We questioned whether extracellular dopamine metabolism may have dampened the effect of the reuptake blocker, thus diminishing the predicted increase in endogenous dopamine levels. Therefore, we repeated this experiment in the presence of bifemelane, a monoamine oxidase A & B inhibitor. Under these conditions, we found further reductions in spontaneous activity when DAT was blocked, as indicated by a significantly reduced response ratio (Fig. 5Bi; n = 6, one-way ANOVA, F_(2,19)_ = 18.0, p < 0.001). Burst analysis revealed significantly fewer episodes (Fig. 5Bii; n = 6, two-way ANOVA, F_(2,20)_ = 11.8, p = 0.0004) with no change in amplitude (Fig. 5Biii; n = 6, two-way ANOVA, F_(2,20)_ = 1.3, p = 0.3). We followed these experiments up with high-performance liquid chromatography (HPLC) to verify endogenous levels of dopamine. In P3 lumbar spinal cords (n = 11) we detected low levels of dopamine; in P60 adults (n = 17) we detected a threefold increase in dopamine levels (Fig. 5Biv; Mann–Whitney U = 10.0, T = 76, p < 0.001). Thus, our in vitro experiments indicated that low levels of endogenous dopamine play a role in D_2_-mediated inhibition.

**Figure 5:**
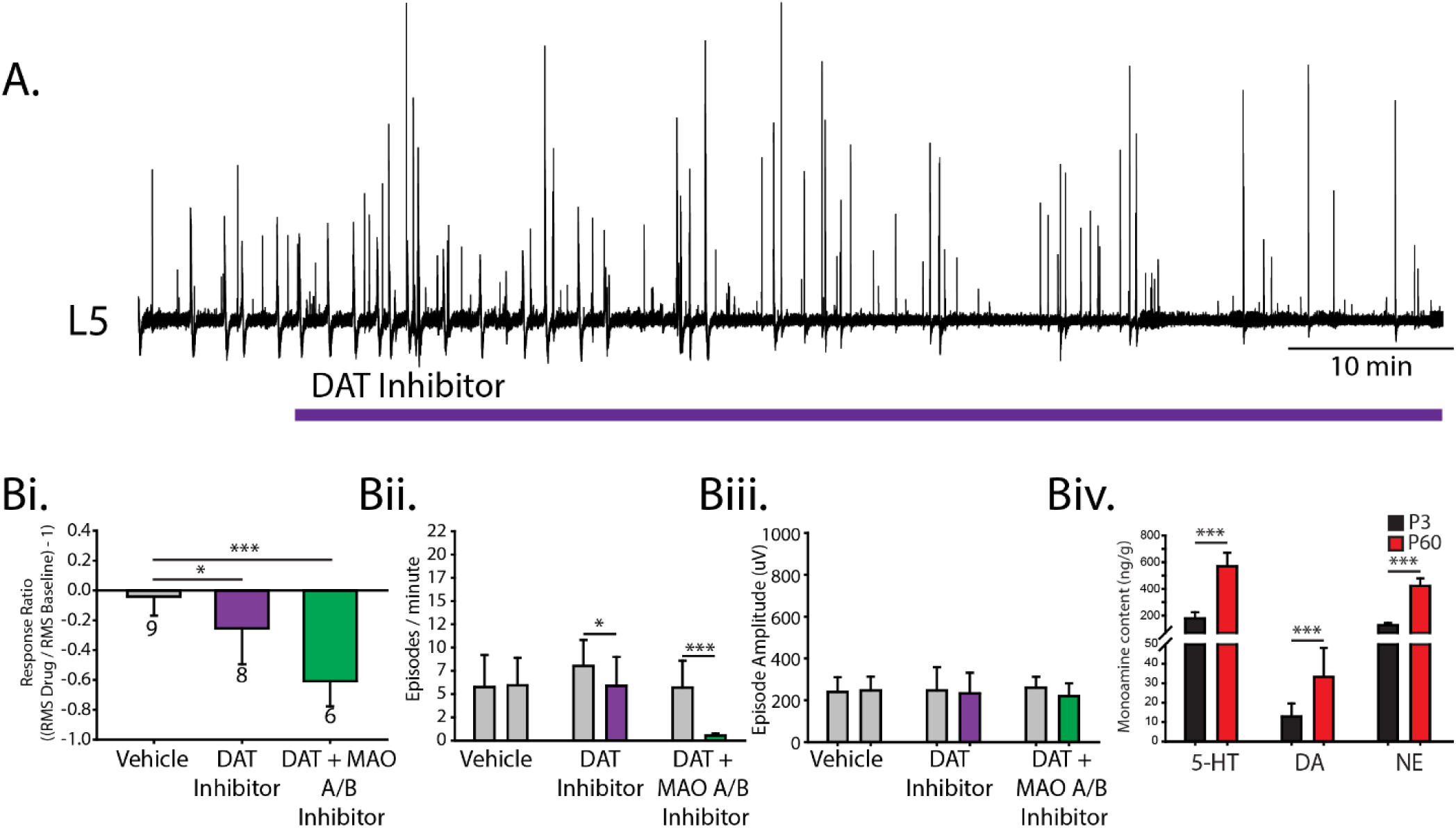
Inhibitory actions of endogenous dopamine in the neonatal mouse spinal cord. A. Single ventral root (L5) neurogram of spontaneous activity after blocking the dopamine transporter (DAT) with the DAT antagonist GBR 12909 (purple) to increase the endogenous dopamine level. Dopamine reuptake was also blocked in the presence of monoamine oxidase A & B inhibitor bifemelane (green) to reduce dopamine metabolism. Bi illustrates the response ratio; negative and positive values indicate inhibition and excitation, respectively. The number below each bar represents the number of preparations for each experiment. Spontaneous episode occurrences per minute (Bii) and amplitude (Biii) were measured within the epochs where the response ratio was calculated. Histograms present mean values ± SD and asterisks indicate a significance level of post hoc analyses (*p < 0.05, **p < 0.01, ***p < 0.001) following one-way ANOVA for response ratios, and two-way ANOVA for burst occurrences and amplitude that compared between conditions. Biv. Endogenous levels of serotonin (5-HT), dopamine (DA) and norepinephrine (NE) were measured in neonatal (P3) and adult (P60) lumbar spinal cords with high-performance liquid chromatography.

### D_1_ receptor activation increases motoneuron excitability by reducing afterhyperpolarization properties

In the next set of experiments, we were interested in determining the cellular mechanisms that mediate dopamine’s complex modulatory effects on spinal network output. As integrators of premotor network activity that generate many of the rhythmic outputs of the spinal cord, motoneurons are ideally suited to amplify spontaneous activity and respond to dopaminergic modulation. Given that they not only serve as the final output for spinal networks, but they also participate in the generation of rhythmic activity [40], we initially selected motoneurons as a locus for determining the cellular mechanisms for dopaminergic excitation and inhibition. We made whole-cell patch-clamp recordings from 75 motoneurons (across 42 animals). Some of these motoneurons (n = 18 across seven animals) were filled with fluorescein-conjugated dextran amine (FITC; Molecular Probes, Inc., Eugene Oregon) and verified post hoc using immunohistochemistry for the presence of choline acetyltransferase (ChAT; Fig. 6A). One hundred percent of filled cells were ChAT positive indicating that we were indeed recording from motoneurons. Electrical properties measured in a group of these cells (n = 8 cells across five animals) were comparable to cells from putative motoneurons recorded in the remaining experiments (Table 1) indicating that these cells are also likely motoneurons. Motoneurons also display characteristic time-dependent changes in repetitive firing frequency during sustained depolarizing current injections (Fig. 6 Cii) [41–43]. We found that all but two cells demonstrated a time-dependent reduction in firing rate throughout the duration of a 0.5 s depolarizing current injection, as indicated by the ratio between maximum steady-state and first spike interval frequencies. This phenomenon is known as spike frequency adaptation (SFA; mean SFA ratio = 0.47 ± 0.19). In addition, doublet firing occurred at the onset of repetitive firing in 5 cells (8%).

**Table 1:** Baseline motoneuron intrinsic properties. Passive, spike, and repetitive firing properties of motoneurons with dopamine or dopamine agonists did not differ at baseline in this series of experiments. Motoneuron identity was verified in a subset of experiments where cells were filled with fluorescein and verified post hoc for expression of choline acetyltransferase (ChAT); 100% of cells were ChAT^+^ indicating that they were indeed motoneurons. Data are means ± SD and analyzed using one-way ANOVAs for each property.

| **Motoneuron basal properties (N= 65 MNs, 40 animals (P0-P4))** |  |  |  |  |  |  |  |  |  |
| --- | --- | --- | --- | --- | --- | --- | --- | --- | --- |
| **Property** | **Chat<sup>+</sup> filled MNs (8,5)<sup>1</sup>** | **Vehicle<sup>2</sup> (12,6)<sup>2</sup>** | **1 μM DA (7,4)<sup>3</sup>** | **5 μM DA (7,4)<sup>4</sup>** | **10 μM DA (6,5)<sup>5</sup>** | **100 μM DA (5,4)<sup>6</sup>** | **SKF 81297 (8,5)<sup>7</sup>** | **Quinpirole (12, 7)<sup>8</sup>** | **(7,56) F, p** |
| **Passive properties** |  |  |  |  |  |  |  |  |  |
| Capacitance (pF) | 203±54 | 203±35 | 193±55 | 194.3±45 | 149±66 | 244±54 | 199±50 | 210±43 | 1.6, 0.15 |
| V<sub>m</sub> (mV) | -73±4.1 | -74±4.3 | -75±4.1 | -73±4.7 | -73±3.7 | -76±3.5 | -72±6.1 | -73±5.0 | 0.7, 0.7 |
| I<sub>Holding -75mV</sub> (pA) | -86±148 | -25±138 | 26±125 | -63±85 | 23±150 | -101±99 | -38±87 | -74±62 | 1.16, 0.3 |
| R<sub>in</sub> (MΩ) | 50±15 | 48±24 | 59±44 | 70±83 | 115±84 | 40±9.2 | 71±28 | 49±14 | 2.0, 0.07 |
| Rheobase (pA) | 678±225 | 747±386 | 596±367 | 464±286 | 443±390 | 762±318 | 580±298 | 633±260 | 0.9, 0.5 |
| **Spike Properties** |  |  |  |  |  |  |  |  |  |
| AP<sub>TH</sub> (mV) | -51±4.4 | -55±4.4 | -55±4.2 | -52±5.9 | -56±3.5 | -53±2.5 | -54±4.3 | -55±4.4 | 1.0, 0.4 |
| AP<sub>AMP</sub> (mV) | 66±5.0 | 71±5.9 | 75±6.9 | 76±13 | 76±4.2 | 69±5.7 | 71±9.6 | 72±6.1 | 1.6, 0.2 |
| AP<sub>rise time</sub> (ms) | 0.45±0.2 | 0.33±0.1 | 0.37±0.1 | 0.4±0.1 | 0.44±0.1 | 0.32±0.04 | 0.35±0.05 | 0.33±0.1 | 1.6, 0.2 |
| AP<sub>half width</sub> (ms) | 0.59±0.15 | 0.60±0.1 | 0.64±0.2 | 0.78±0.3 | 0.75±0.2 | 0.62±0.07 | 0.68±0.1 | 0.68±0.19 | 1.2, 0.3 |
| AHP<sub>duration</sub> (ms) | 73.7±29 | 70±18 | 88±34 | 76±13 | 108±38 | 74±16 | 69±23 | 76±26 | 1.5, 0.2 |
| AHP<sub>AMP</sub> (mV) | -3.9±1.1 | 3.6±2.2 | 3.1±1.4 | 3.2±1.4 | 3.4±2.1 | 4.7±1.2 | -4.2±1.9 | -4.3±1.4 | 0.8, 0.6 |
| **Repetitive Firing Properties** |  |  |  |  |  |  |  |  |  |
| SS F/I Slope (Hz/pA) | 0.053±0.01 | 0.044±0.008 | 0.058±0.008 | 0.052±0.018 | 0.053±0.03 | 0.051±0.007 | 0.046±0.008 | 0.053±0.012 | 1.2, 0.3 |
| Repetitive firing TH (pA) | 605±446 | 498±254 | 410±271 | 354±200 | 338±327 | 515±253 | 367±186 | 494±173 | 0.8, 0.6 |
| Max SS firing rate (Hz) | 59±8.1 | 59±8.3 | 51±13 | 46±7.9 | 52±8.9 | 55±12 | 57±9.5 | 58±12 | 1.9, 0.09 |
| First spike latency (ms) | 146±86 | 116±57 | 124±77 | 91±34 | 113±79 | 136±55 | 132±67 | 142±50 | 0.5, 0.8 |

**Figure 6:**
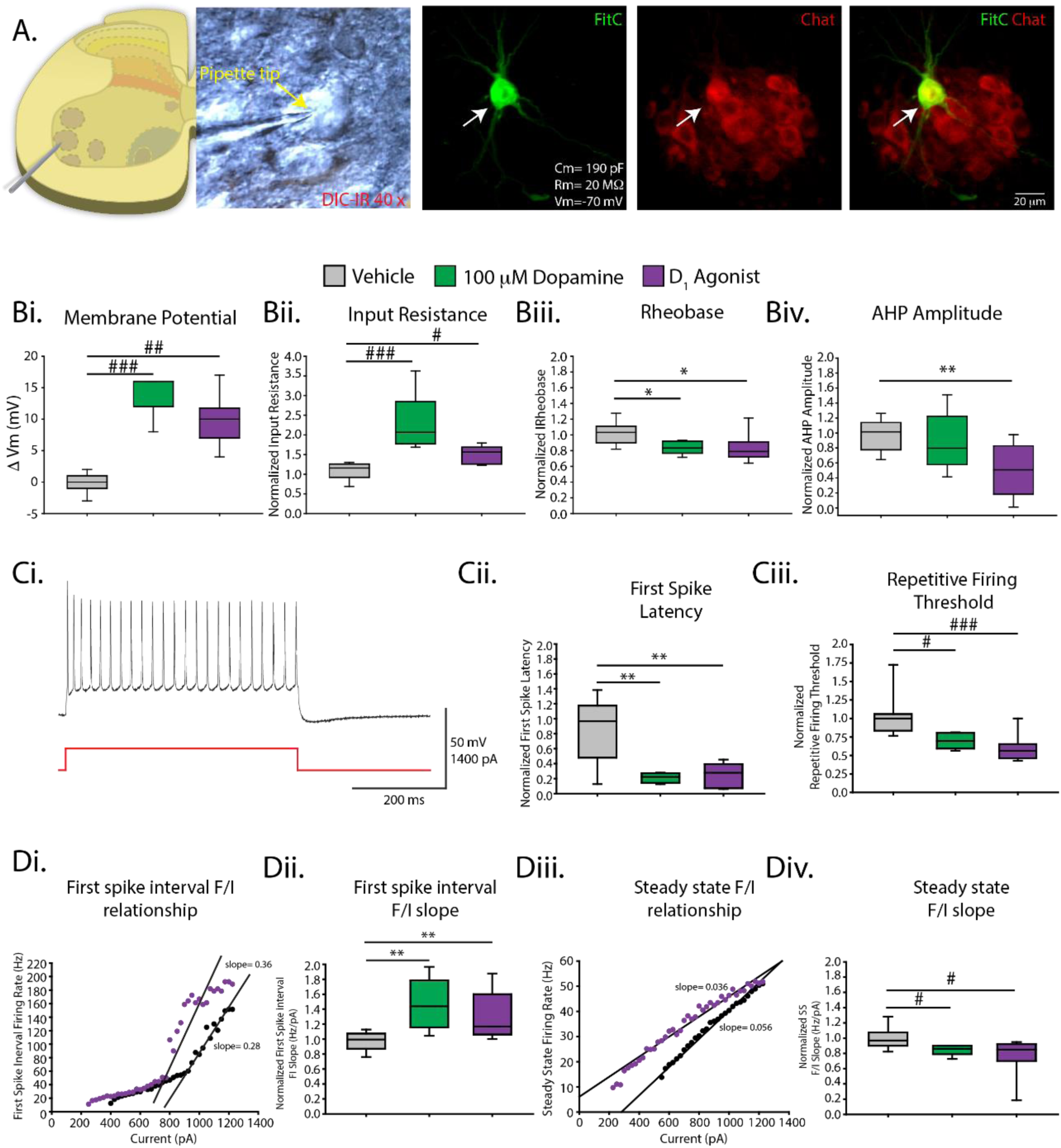
High concentrations of dopamine and D_1_ agonists increase motoneuron excitability. A. Whole-cell patch-clamp recordings were obtained from motoneurons visualized with infrared differential interference contrast (IR-DIC) in lumbar slices of neonatal (P0–P4) mice. A subset of cells were filled with fluorescein (FITC, green) and verified post hoc with immunohistochemistry for choline acetyltransferase (ChAT, red). B–D. High concentrations of dopamine (100 µM, green) and the D_1_ agonist SKF 81297 (20 µM, purple) increased motoneuron excitability by modulating various passive, spike, and repetitive firing properties. All measures were compared to that of time-matched vehicle control (grey). Ci. A typical motoneuron trace following current injection. Di, Diii display representative frequency–current (FI) plots from a single cell before (black dots) and after (purple dots) application of the D_1_ agonist. Box and whisker plots display interquartile range (boxes), median (horizontal black lines), max, and minimum value in data range (whiskers). Asterisks denote significance level of post hoc tests (*p < 0.05, **p < 0.01, ***p < 0.001) following one-way ANOVA.

Consistent with previous reports from our laboratory [44,45] and with our network recordings in this report, 100 µM of dopamine increased motoneuron excitability (n = 5 cells across four animals); we reproduced this effect with the D_1_ agonist SKF 81297 (20 µM; n = 8 cells across five animals). Both 100 µM dopamine and the D_1_ agonist depolarized the membrane potential (Fig. 6Bi; H_(2)_ = 18.9, p < 0.001), increased negative holding current (vehicle, −7.1 ± 33 pA; DA, −247 ± 78 pA; D_1_, −174 ± 49 pA; H_(2)_ = 18.9, p = 0.001) and input resistance (Fig. 6Bii; H_(2)_ = 16.0, p < 0.001), and decreased rheobase (Fig. 6Bii; F_(2,22)_ = 5.0, p = 0.016) beyond that of the time-matched vehicle control. We found no change in spike rise time (F_(2,22)_ = 1.0, p = 0.4) or half width (F_(2,22)_ = 0.8, p = 0.5). The D_1_ agonist reduced the amplitude of the afterhyperpolarization (AHP); however, 100 µM dopamine did not reduce AHP amplitude beyond that of a time-matched vehicle control (Fig 6Biv; F_(2,22)_ = 7.7, p = 0.003). Frequency–current (FI) relationships were measured for steady state and first spike interval during a series of depolarizing current pulses (Fig. 6Ci). Both 100 µM dopamine and the D_1_ agonist reduced the latency to first spike beyond that of the time-matched vehicle control (Fig. 6Cii; F_(2,17)_ = 9.6, p = 0.002). Dopamine (100 µM) and the D_1_ agonist increased the slope of the exponential region of the FI relationship for the first spike interval (Fig. 6Di–Dii; F_(2,22)_ = 8.4, p = 0.002) and reduced the slope of the steady-state FI relationship (Fig. 6Diii–Div; H_(2)_ = 9.4, p = 0.009). The reduction in steady-state slope was due to a leftward shift in steady-state FI relationship (Fig. 6Diii) characterized by a reduction in the threshold for repetitive firing (Fig. 6Ciii; H_(2)_ = 14, p < 0.001) with no change in the maximum steady-state firing rate (H_(2)_ = 1.4, p = 0.5). These results indicate that activation of D_1_ receptors elicit consistent effects as high concentrations of dopamine on motoneuron excitability and is a likely mechanism contributing to dopaminergic excitation of motor output.

### Dopaminergic inhibition through D_2_ - receptor hyperpolarization of distributed populations of ventral interneurons

We next set out to determine the cellular mechanisms that mediate the inhibitory effects of dopamine on spinal network output with motoneurons as our first target. In contrast to our network recordings, low concentrations of dopamine (1–10 µM; n = 20 cells across 14 animals; Table 1) and the D_2_ agonist quinpirole (20 µM; n = 12 cells across seven animals; Table 1) did not alter any passive, spike or repetitive firing properties of motoneurons beyond that of the time-matched vehicle control (n = 12 cells across six animals). A lack of responsiveness was not due to the diversity of motoneurons as there was no correlation between cell capacitance and changes in membrane potential or holding current elicited by low concentrations of dopamine (r = −0.3, p = 0.13) or quinpirole (r = −0.1, p = 0.72). These results suggest that the inhibitory actions of dopamine on spinal network output are not due to D_2_-receptor inhibition of intrinsic motoneuron excitability; instead, dopamine may be acting on premotor interneurons.

Many of the premotor interneurons that produce the rhythmic activities generated by the spinal cord are distributed across lamina’s VII - X of the ventral lumbar spinal cord. We next recorded from ventral interneurons located in lamina VII–X to determine a cellular locus for D_2_-mediated inhibition of spinal network output (n = 30 cells across 17 animals; Table 2). Quinpirole produced a sustained hyperpolarization of the resting membrane potential in 33% of interneurons (Fig. 7B, C, Di; n = 10; dVm = −4.8 ± 2.4 mV; two-way ANOVA, F_(2,37)_ = 26.1, p < 0.0001) and a transient hyperpolarization of membrane potential in 10% of interneurons (n = 3; dVm = −5.16 ± 1.9 mV) that persisted for 209 **±** 108 s before returning to baseline levels. The change in resting membrane potential was greater in the responders compared to non-responders (dVm responders = - 4.8 ± 2.4 mV; dVm non-responders = 1.6 ± 1.9 mV; unpaired t-test t_(28)_ = 7.8, p < 0.0001). Quinpirole reduced the input resistance (Fig. 7Dii; two-way ANOVA, F_(2,37)_ = 4.1, p = 0.025) and increased the spike rise time (Fig. 7Div; two-way ANOVA, F_(2,35)_ = 5.2, p = 0.01; Dunn post hoc test, p=0.02) but not the half width (F_(2,35)_ = 2.6, p = 0.086) in the group of responding interneurons (“responders”). For both responders and non - responders, quinpirole did not alter rheobase (Fig. 7Diii; two-way ANOVA, F_(2,37)_ = 0.5, p = 0.6), action potential threshold (two-way ANOVA, F_(2,37)_ = 0.3, p = 0.7), AHP amplitude (two-way ANOVA, F_(2,37)_ = 0.04, p = 0.96), duration (two-way ANOVA, F_(2,37)_ = 0.8, p = 0.45), threshold for repetitive firing (two-way ANOVA, F_(2,37)_ = 0.01, p = 0.98), maximum steady-state firing rate (Fig. 7Dvi; two-way ANOVA, F_(3,37)_ = 0.8, p = 0.5) or steady-state FI slope (Fig. 7Dv: two-way ANOVA, F_(2,37)_ = 0.9, p = 0.4). The capacitance (two-way ANOVA, F_(2,37)_ = 4.2, p = 0.02) and holding current (two-way ANOVA, F_(2,37)_ = 4.9, p = 0.02) were higher in responders than in non - responders (Table 2) and a greater proportion of responders were localized to more medial regions of spinal slices (Fig. 7B), where putative commissural interneurons reside [46].

**Table 2:** Baseline ventral interneuron intrinsic properties. Passive, spike, and repetitive firing properties of ventral interneurons that responded to quinpirole with sustained hyperpolarization of membrane potential (responders) were compared with those that did not respond (non-responders) and also retrogradely labelled descending commissural interneurons (dCINs). Responders had higher capacitance and holding current than non-responders and dCINs. Data are means ± SD; F and p values are reported from one-way ANOVAs for each property. Red values highlight variables where significant main effects were detected with p < 0.05. Superscript numbers reflect significant differences between respective conditions from post hoc analysis.

| **Ventral Interneurons (n=40 cells, 22 animals (P0-4))** |  |  |  |  |
| --- | --- | --- | --- | --- |
| **Property** | **Non-responders<sup>1</sup>**<br>**(n=20,8)** | **Responders<sup>2</sup>**<br>**(n=10,9)** | **dCINs<sup>3</sup>**<br>**(n=10,5)** | **(2,37) F, p** |
| **Passive Properties** |  |  |  |  |
| Capacitance (pF) | 47±15<sup>2</sup> | 73±40<sup>1</sup> | 46±17<sup>2</sup> | 4.2, 0.02 |
| V<sub>m</sub> (mV) | -66±4.3 | -66±5.2 | -66±6.0 | 0.02, 0.98 |
| I<sub>Holding -75mV</sub> (pA) | -10.8±16<sup>2</sup> | -57±52<sup>1</sup> | -26±18 | 4.9, 0.02 |
| R<sub>in</sub> (MΩ) | 759±466 | 497±248 | 719±326 | 1.6, 0.1 |
| Rheobase (pA) | 84±103 | 59±13 | 47±13 | 0.9, 0.4 |
| **Spike Properties** |  |  |  |  |
| AP<sub>TH</sub> (mV) | -55±4.6 | -55±2.9 | -56±2.3 | 0.2, 0.8 |
| AP<sub>AMP</sub> (mV) | 71±9.3 | 71±8.2 | 72±7.3 | 0.05, 0.95 |
| AP<sub>rise time</sub> (ms) | 0.99±0.48<sup>3</sup> | 0.92±0.3<sup>3</sup> | 1.4±0.25<sup>1,2</sup> | 4.2, 0.023 |
| AP<sub>half width</sub> (ms) | 1.2±0.5 | 1.4±0.6<sup>3</sup> | 1.9±0.4<sup>2</sup> | 6.2, 0.005 |
| AHP<sub>duration</sub> (ms) | 173±96 | 193±76 | 172±81 | 0.2, 0.8 |
| AHP<sub>AMP</sub> (mV) | -4.9±2.7 | -4.8±2.1 | -4.8±2.7 | 0.02, 0.98 |
| **Repetitive Firing Properties** |  |  |  |  |
| SS F/I slope (Hz/pA) | 0.24±0.07 | 0.21±0.09 | 0.2±0.1 | 1.0, 0.4 |
| Repetitive firing TH (pA) | 26±14 | 31±12 | 28±13 | 0.5, 0.6 |
| Max firing rate (Hz) | 43±16.3 | 39±11.6 | 32±5.9 | 2.1, 0.13 |
| Fist spike latency (ms) | 143±81 | 146±95 | 117±87 | 0.45, 0.6 |

**Figure 7:**
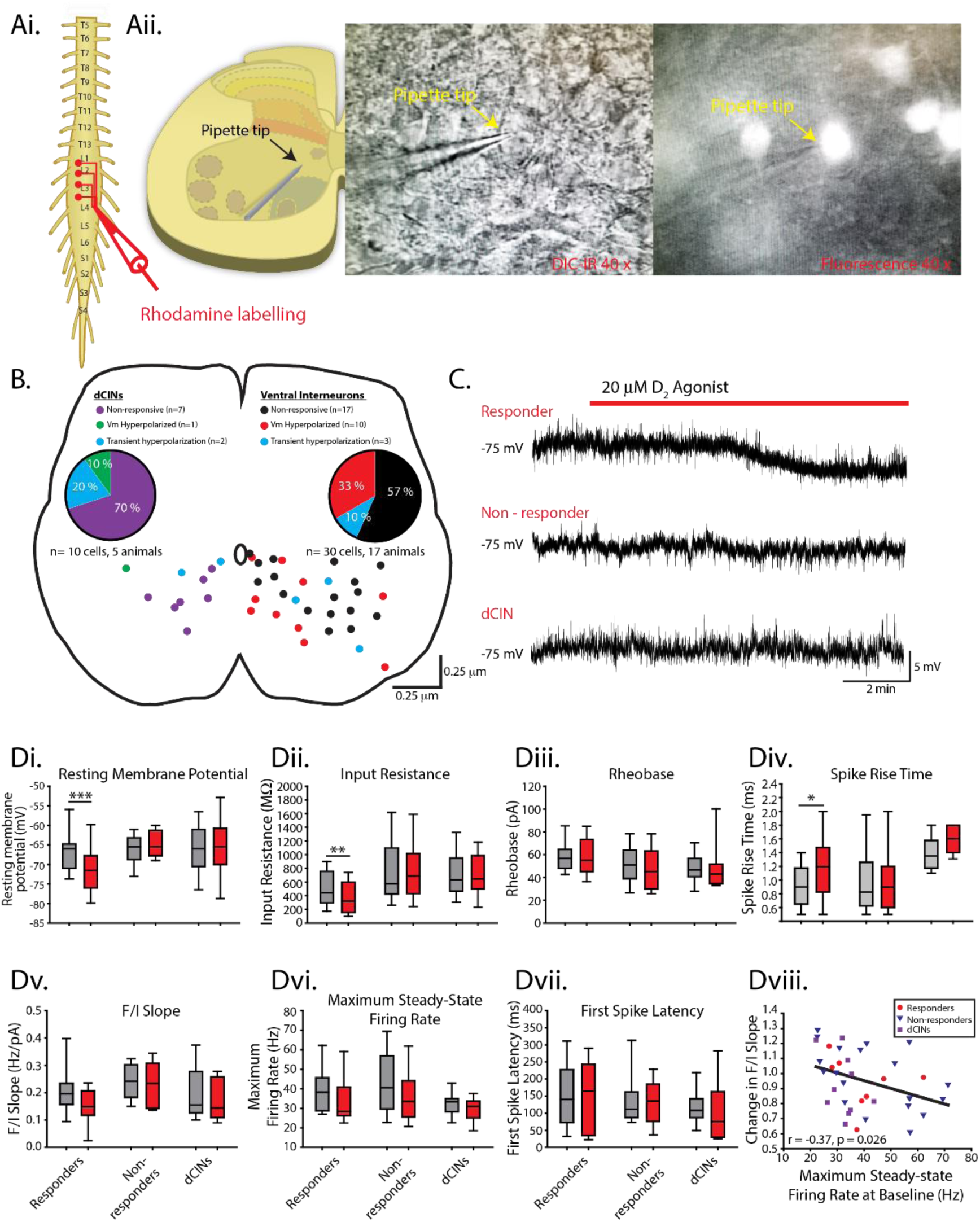
D_2_ agonists hyperpolarize a proportion of ventral interneurons. Whole-cell recordings obtained from lamina VII–X interneurons. Ai. Descending commissural interneurons (dCINs) were retrogradely labelled with rhodamine-conjugated dextran amine crystals inserted into the ventrolateral funiculus at L4 level. Aii. Cells were visualized in transverse lumbar slices under infrared differential interference contrast (IR-DIC) or epifluorescent illumination. B. Ventral interneuron location was measured relative to the central canal and X-Y positions normalized to the distance of ventral and lateral borders of lumbar slices. C. Whole-cell current-clamp recordings depict responsiveness of interneuron resting membrane potential to the D_2_ agonist (quinpirole, 20 µM) in responders, non - responders, and dCINs. A proportion of ventral interneurons responded to quinpirole with either a sustained (B, red and green dots) or transient (B, blue dots) hyperpolarization of the membrane potential. A reduction in input resistance and increase in spike rise time accompanied hyperpolarization in sustained responders …. … (Dii, Div) Box and whisker plots display interquartile range (boxes), median (horizontal black lines), max, and minimum value in data range (whiskers). asterisks denote significance level from post hoc tests (*p < 0.05, **p < 0.01, ***p < 0.001) following two-way ANOVA. Dviii. Reductions in steady-state frequency-current (FI) slope of all cells following application of quinpirole was greatest in cells that had a higher maximum steady-state firing rate at baseline.

We next set out to determine the type of interneurons that were hyperpolarized by quinpirole. Given that many of the responding cells were located medially, we next targeted descending commissural interneurons (dCINs; n = 10 cells across five animals; Table 2) since this population can be identified based on anatomical connectivity [47,48], display intrinsic burst properties [49] and are rhythmically-active during neurochemically-evoked fictive locomotion [50]. dCINs were retrogradely labelled with tetramethylrhodamine-conjugated dextran amine (molecular weight (MW) 3000; Molecular Probes, Inc.) inserted into the ventrolateral funiculus at the L4 segment (Fig. 7A). In contrast to our hypothesis, only one dCIN responded with a sustained hyperpolarization and two were transiently hyperpolarized (Fig. 7B) by quinpirole. Quinpirole did not alter any passive, spike, or repetitive firing properties of dCINs (n = 10; Fig. 7D). These data suggest that the dCINs, although responsive in similar proportions to our global interneuron survey, do not exclusively account for the 33% responding group and, therefore, are likely, not responsible for the observed network effects. Interestingly, when all interneuron data were pooled (n = 40 cells), irrespective of responsiveness as indicated by changes in resting membrane potential, quinpirole had the greatest effect in cells that had a higher maximum steady-state firing rate at baseline (Fig. 7DViii; r = −0.37, p = 0.026). There was no correlation between baseline FI slope and changes in FI slope in response to quinpirole (r = −0.09, p = 0.6). While dCINs can be identified anatomically, they are heterogeneous with respect to their neurotransmitter phenotype [51] and as a result have varying contributions to network activities [46,50,52].

We therefore next targeted V3 interneurons which are exclusively glutamatergic, contribute to the stabilization of locomotor-like rhythmicity and can be identified genetically based on the expression of the Sim1 transcription factor [53].

Given that dopamine inhibits ventral root-evoked locomotor activity, which may be mediated by this circuit [56], through D_2_-receptor signalling [31], we hypothesized that V3 interneurons may be a cellular locus for D_2_-mediated inhibition of spinal network activity.

Consistent with our global interneuron survey, quinpirole produced a sustained hyperpolarization of the resting membrane potential in a proportion of V3 interneurons (n=5 cells; 27%) that we recorded from (total V3 interneurons n=23 cells, 7 animals) and transient hyperpolarization in 3 (13%) V3 interneurons. The magnitude of the response in the 5 cells that responded with a sustained hyperpolarization was variable and approached, but did not reach, significance (Supp Figure 1; paired t-test: t_(4)_= 2.5, p = 0.06); however, did reach significance when the cells that responded with a transient hyperpolarization were included in the analysis (paired t-test: t_(6)_ = 2.7, p = 0.03). Consistent with our global interneuron survey the change in resting membrane potential elicited by quinpirole was significantly greater in responding (n = 5; dVm = −1.9 ± 1.9 mV) compared to non-responding (n = 15; dVm = −0.3 ± 1.0 mV) V3 interneurons (unpaired t-test: t_(16)_ = 3.8, p = 0.002).

## Discussion

Dopamine is a monoamine neuromodulator that is important for the control of rhythmically active motor circuits across phyla (reviewed by [8]) but is probably best known in vertebrates for the control of dedicated circuits in the basal ganglia that control action selection (reviewed by [57]). Work in small circuits of invertebrates has established that circuit connectomes define the constraints on which networks operate and that neuromodulators diversify outputs by altering intrinsic and synaptic properties of the neurons that compose the circuit (for reviews see [1–3]. In line with this, the distribution of receptors within circuits constrain the effect of neuromodulators on circuit output. For example, dopamine is exclusively inhibitory in spinal circuits of *Xenopus* tadpoles prior to free-swimming stages [58] due to expression of D_2_ but not D_1_ receptors. We show that dopamine has has bidirectional concentration-dependent effects on spinal network output in neonatal mice where all dopamine receptor types are expressed which is consistent with what has been reported in tadpoles at free swimming-stages [12]. Our data highlights that neuromodulator concentration is also important because receptors have varying ligand affinities which underlie concentration-dependent actions of modulators. Although dopamine predominantly inhibits spinal output in neonatal mice, similar to pre-free-swimming tadpoles, it is primarily due to the concentration of spinal dopamine, not the distribution of receptors. Our previous work shows that neuromodulation of mammalian spinal networks is dependent on network excitability state [13] which is consistent with findings from invertebrates [14,15]. Our current work shows that receptor mechanisms and concentration-dependent control of spinal network output is also state-dependent. This is important because receptor expression, modulator concentration and network excitability are not fixed and fluctuate dynamically [59,60]. Therefore, these three factors need to be considered if we wish to understand how networks create diverse neuromodulator-dependent outputs (Figure 8A).

**Figure 8:**
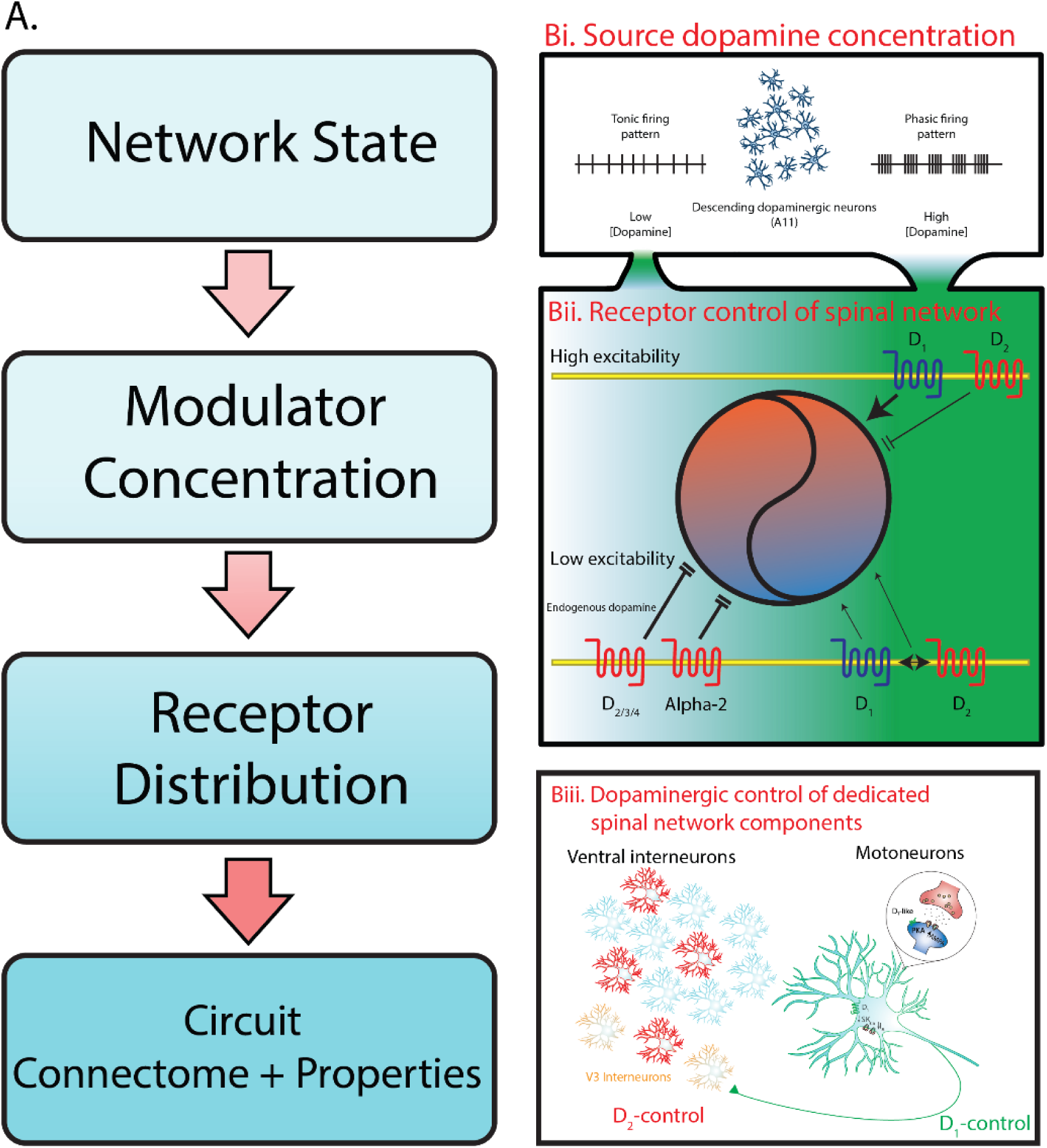
Dopamine exerts state and receptor-dependent control of spinal motor networks. A. Neural network output is dependent on multiple factors. Circuit connectomes and intrinsic properties defining constraints of a network output. Similarly, the distribution of neuromodulator receptors defines the constraints on which neuromodulators can alter these properties. Varying ligand affinities of these receptors determine dose-dependent effect of neuromodulators and network state dictates the dose-dependent effects of neuromodulators. Bi. Previous work has demonstrated that tonic and phasic firing patterns of dopamine cells lead to the release of low and high concentrations of dopamine, respectively [10]. Bii. During a low excitability state (blue part of the circle), low concentrations of dopamine (denoted by the green gradient) acted on D_2_, D_3_, D_4_ and α2-adrenergic receptors in parallel to inhibit (parallel stop lines) spinal motor output. As dopamine levels rise, activation of D_1_ and co-activation of D_2_ receptors increase (arrowheads) spinal network output. During a high excitability state (orange part of the circle), the actions of low dopamine are not apparent and at higher concentrations, activation of D_1_ receptors boost … … spinal network output and activation of D_2_ receptors slows rhythmic activity [13]. Biii. Higher concentrations of dopamine act through D_1_ receptors to increase motoneuron excitability by reducing A-type and SK_Ca_-dependent calcium conductance. D_2_-mediated inhibition of spinal network output is triggered by reduced excitability in a proportion of ventral interneurons (red cells).

### A physiologically silent excitatory D_1_ pathway?

Previous work has shown predominantly excitatory D_1_-receptor mediated effects of dopamine on fictive locomotion which is characteristic of the network operating in a high excitability state [13] - albeit in these studies, higher concentrations of dopamine are necessary to elicit observable effects [20–25]. Here, we show that endogenous levels of dopamine within the spinal cord of neonatal mice are low. Even though HPLC suggests low concentrations of dopamine in the neonatal spinal cord, a critical question that we addressed is what would happen when we manipulated endogenous dopamine? We accomplished this by blocking dopamine reuptake and metabolism and found that the effects were not excitatory, but inhibitory.

Based on this, we suggest that although present, the excitatory D_1_-mediated pathway is ‘physiologically silent’ during early postnatal stages given that endogenous levels are not sufficient to activate this pathway. A similar phenomenon has been reported for glutamatergic synapses in developing circuits of the hippocampus which express NMDA but not AMPA receptors. As a result, glutamatergic synapses fall ‘physiologically silent’ as the release of glutamate does not produce sufficient depolarization of the postsynaptic membrane to remove the magnesium block from the pore of the NMDA channel [61,62].

Excitability state also influences the receptor mechanisms and therefore concentration-dependent control of modulators on network output. In the spinal cord, D_1_ pathways may be more important for the regulation of spinal circuits operating in higher excitability states. For example, fictive locomotor rhythms are drastically impacted when D_1_ receptors are manipulated whereas manipulation of D_2_ receptors elicit only subtle changes in rhythm frequency, with rhythm robustness being maintained [23] (Figure 8Bii.). In the adult animal; however, network excitability increases, and spinal dopamine levels are higher due to increases in descending inputs. In line with this, optogenetic activation of the dopaminergic A11 leads to an increase in motor activity [63]. Similarly, the excitatory D_1_ system is more important for the control of stepping movements whereas the inhibitory D_2_ system plays less of a role [22,64]. Instead, the inhibitory D_2_ pathway may be more important in maintaining network quiescence during periods of immobility such as has been inferred network excitability are low, that inhibition of motor output prevails (Figure 8Bii.).

Thus, this points to the receptor-specific and therefore concentration-dependent control of modulators on network output being strongly influenced by the state of the network on which they are acting.

### Dedicated network components segregate excitatory and inhibitory control of spinal networks

Dedicated circuits regulated by non-overlapping populations of neurons that express D_1_ and D_2_ receptors compose the direct and indirect pathways of the basal ganglia in vertebrates and have also been reported in the superior colliculus of rodents [65]. Our work suggests dedicated network elements within the spinal cord that are regulated by D_1_ and D_2_ pathways (Figure 8Biii.). Specifically, we found that D_1_ receptors excite motoneurons through similar mechanisms that have been previously reported [44,45], but are not affected by low concentrations of dopamine or D_2_ agonists. This points to the possibility of a dedicated D_1_-dependent circuit that could underlie the generation of rhythmic activities elicited by high concentrations of dopamine. Motoneurons compose key rhythm generating elements in invertebrate circuits [66–68] and also participate in rhythm generation in vertebrates [69], including rodents [32,40,56,70,71]. V3 interneurons are one subclass of genetically-defined spinal interneuron that are important for the generation of rhythmic activities in mammalian spinal networks [53,55,72] and receive recurrent excitatory collaterals from motoneurons in rodents [32]. Motoneurons in the rodent spinal cord also form glutamatergic synaptic connections amongst each other [71] and activation of D_1_ receptors could serve to synchronize motor pools. Previous results demonstrating D_1_- and not D_2_-mediated increases in AMPA conductances on motoneurons [45] support this possibility. We cannot rule out the possibility that there is a degree of overlap between D_1_ and D_2_ controlled network elements within the spinal cord. While we did not examine the D_1_ control of ventral interneurons, we did find that D_2_ receptors hyperpolarize a subset of V3 interneurons. This population is therefore a potential cellular locus where cooperative excitatory D_1_-D_2_ interactions that we report here could occur. D_1_ and D_2_ receptors have been reported in the brain to become co-activated or form heterodimeric complexes that augment neuronal excitability through PLC-dependent increases in intracellular calcium [35–38]. Our pharmacological and immunoprecipitation data indicate that this may also occur in the neonatal mouse spinal cord. Although this pathway may be physiologically silent during early postnatal development, increasing dopamine concentrations at later stages may activate this pathway.

The inhibitory D_2_ pathway on the other hand does not appear to act through the modulation of motoneuron intrinsic or synaptic properties [45] but instead hyperpolarizes a proportion of ventral interneurons. Based on our responsiveness criteria, a majority of cells that responded to a D_2_ agonist with sustained hyperpolarization of the membrane potential were localized more medially in lumbar slices. We tested the hypothesis that a subpopulation of interneurons was D_2_ sensitive and tested both dCINs and genetically identified V3 interneurons. Our hypothesis was not supported, and similar proportions of D_2_ sensitive neurons were found; however, the data suggest that there is heterogeneity in the responsiveness to neuromodulators within a class of genetically defined interneurons and that D_2_ actions may not be localized to a particular class of interneurons.

One possibility is that neuromodulators elicit a robust effect on network output through distributed control across multiple classes of genetically-defined interneurons which would be consistent with findings that the locomotor rhythm generator is also distributed across several classes of interneurons [73–75]. Our network data suggests that low concentrations of dopamine inhibit network output by acting in parallel on D_2_, D_3_, D_4_ and α_2_ receptors. We may therefore have underestimated the cellular targets that underlie dopamine’s robust inhibitory effect on the network given that in this series of experiments we looked at the activation of D_2_ receptors alone.

### Developmental considerations for the endogenous dopaminergic system

This robust inhibitory system may act as a brake on network activity during perinatal development when chloride-mediated synaptic transmission still causes partial depolarization [76]. An inhibitory brake would prevent runaway excitation of spinal network activity during this critical period of development. A robust background inhibitory system mediated by the parallel inhibitory action of D_2_, D_3_, D_4_, and α2-adrenergic receptors activated by low levels of endogenous dopamine would counteract depolarizing chloride-mediated transmission.

Signalling through [76] D_2_-like receptors may also play a role in driving the maturation of spinal networks. In larval zebrafish, D_4_ receptors drive the maturation of spinal locomotor network organization [77] and function leading to changes in locomotor behaviour [78]. Similar processes may also occur perinatally in rodents, in that the preferential activation of the D_2_ receptor system may favour intracellular signalling that results in network reorganization. Serotonin receptors have been found to shape network function and inhibitory synaptic transmission during early postnatal days of rodents [79,80]. Dopamine could, therefore, act analogously via the D_2_-system during perinatal development.

### Conclusions

Here we present evidence for an inhibitory physiological role of dopamine in the regulation of developing mammalian spinal networks. We also demonstrate an excitatory D_1_-mediated pathway that acts through excitation of motoneurons, and possibly recurrent excitatory collaterals to CPG neurons, however given that endogenous levels of dopamine are low, propose that this pathway is physiologically silent. These data advance our understanding of how neuromodulators regulate network output in light of dynamically changing modulator concentrations and levels of network excitability.

## Acknowledgments

We would like to acknowledge support from the Whelan Lab and technical support from Dr. Heather Leduc-Pessah and Rebecca Dalgarno on immunoprecipitation experiments and from Gail Rauw on HPLC experiments. Spinalcore software was a kind gift from Prof A. Lev-Tov (Hebrew University).

## Contributions

SAS performed and analyzed whole-cell and network electrophysiology, NEB, performed and analyzed CoIP, GBB performed and analyzed HPLC, JBF performed recordings from V3 interneurons and SAS, CHTK and SEAE performed IHC and imaging experiments. SAS, CJ-X and PJW interpreted results. SAS prepared figures and drafted the manuscript. TT, YZ and PJW contributed to experimental design and procured funding. SAS, CJ-X and PJW conceived and designed the research. All authors reviewed and approved the final version of the manuscript.

## Methods

### Ethical approval & animals

Experiments were performed on male and female neonatal (P0–P4, n = 262) and adult (P50, n = 17) C57BL/6 mice. A subset of experiments were performed on spinal cord slices obtained from neonatal (P0-4, n=7) *Sim1*^*Cre*/+^; *Rosa*^*floxstop26TdTom*^ which were used to visualize and record from V3 interneurons. All procedures performed were approved by the University of Calgary Health Sciences Animal Care Committee University Committee on Laboratory Animals at Dalhousie University.

### Tissue preparation

We anesthetized all animals by hypothermia. Pups were decapitated and eviscerated to expose the vertebral column and rib cage. The isolated vertebrae and rib cage were transferred to a dish lined with a silicone elastomer (Sylgard; DowDuPont, Midland, MI) and perfused with room-temperature (21 - 23°C) carbogenated (95% O_2_ / 5% CO_2_) aCSF (in mM, 4 KCl, 128 NaCl, 1 MgSO_4_, 1.5 CaCl_2_, 0.5 Na_2_HPO_4_, 21 NaHCO_3_, 30 D-glucose; 310–315 mOsm.). We exposed the spinal cord with a ventral laminectomy and isolated it by cutting the nerve roots that connected it to the vertebral column. The isolated spinal cord was then transferred to a recording chamber, perfused with carbogenated aCSF, and placed ventral side up. The bath temperature was gradually increased to 27°C [81]. We let the spinal cords stabilize for 1 hour before performing experiments.

### Spinal cord slice preparation

Following isolation, spinal cords were transected above the tenth thoracic (T10) and below the first sacral (S1) segments and transferred to a slicing chamber. Pre-warmed liquefied 20% gelatin was used to secure cords to an agar (3%) block that was super-glued to the base of a cutting chamber and immersed in ice-cold, carbogenated, high-sucrose slicing aCSF (in mM, 25 NaCl, 188 sucrose, 1.9 KCl, 10 Mg SO_4_, 1.2 Na_2_HPO_4_, 26 NaHCO_3_; 25 D-Glucose; 340 mOsm). Using a vibratome (Leica, Bussloch, Germany) we cut 250-µm-thick lumbar slices, collected and transferred them to a recovery chamber containing regular carbogenated aCSF (see Tissue Preparation) heated to 32_o_C for one hour, then maintained them at room temperature for at least 30 minutes before transferring them to a recording chamber.

### Labelling of descending commissural interneurons (dCINs)

We retrogradely labelled dCINs by inserting tetramethylrhodamine-conjugated dextran amine crystals (MW 3000; Molecular Probes, Inc.) into a cut in the ventrolateral funiculus at the L4 segment. Spinal cords recovered for 4 hours to allow retrograde uptake of the fluorescent dye. Fluorescently-labelled cells were visualized with epifluorescent illumination.

### Electrophysiological recordings

Extracellular neurograms were recorded by drawing ventral roots of the second (L2) and fifth (L5) lumbar segments into tight-fitting suction electrodes fashioned from polyethylene tubing (PE50). Signals were amplified 1000× in total via 10× pre-amplification and 100× second-stage amplification (Cornerstone EX4-400 Quad Differential Amplifier; Dagan Corporation, Minneapolis, MN). Amplified signals were band-pass filtered (0.1–1000 Hz) and digitized at 2.5 kHz (Digidata 1440A/1550B; Molecular Devices, Sunnyvale, CA). Data were acquired in Clampex 10.4/10.7 software (Molecular Devices) and saved on a Dell computer for offline analysis. All experiments were performed on spinal cords naïve to drugs and experimental treatment.

### Whole-cell patch-clamp recordings

Spinal cord slices were gently transferred to a recording chamber perfused with room-temperature carbogenated recording aCSF (in mM, 128 NaCl, 4 KCl, 1.5 CaCl_2_, 1.0 MgSO_4_, 0.5 Na_2_HPO_4_, 21 NaHCO_3_, 30 D-glucose; approximately 310 mOsm) and stabilized in the recording dish with a stainless-steel harp. We gradually heated aCSF to 27°C. Slices were visualized (Olympus BX51WI; Olympus Corporation, Tokyo, Japan) under 5× magnification and putative motoneurons identified using a 40× objective with infrared differential interference contrast (IR-DIC) illumination. We identified putative motoneurons based on their location in the ventrolateral spinal cord and a soma diameter of greater than 20 µm. A cohort of motoneurons was passively filled with fluorescein dextran amine (FITC; MW 3000; 200μM; Molecular Probes, Inc.) added to the intracellular solution to visualize and localize the recorded cells, for 20–60 minutes and motoneuron identity was verified post hoc by performing immunohistochemistry for choline acetyltransferase (ChAT). To verify motoneuron identity in subsequent experiments, basal biophysical properties of putative motoneurons were compared with the FITC-filled ChAT-positive cells. We identified no differences in basal biophysical properties of ChAT-positive identified cells, compared with cells in other experiments identified by position and size (Table 1). In one set of experiments, we recorded from ventral interneurons in lamina VII–X, descending commissural interneurons (dCINs) and V3 interneurons of the lumbar spinal cord (Table 2). We measured the distance of each interneuron from the central canal (LinLab2; Scientifica, Uckfield, UK) and normalized position, relative to lateral and ventral borders, and plotted the normalized positions on a template transverse spinal cord. Responsiveness was determined by the change in resting membrane potential. All experiments were performed on one cell per slice to ensure that all cells were naïve to treatment.

Recording electrodes were pulled from borosilicate capillary tubes (O.D. = 1.5mm, I.D. = 0.86 mm) using a Flaming/Brown Model P-97 micropipette puller (Sutter Instrument, Novato, CA). Pipettes pulled to record from motoneurons, and interneurons were within the range of 3–5 MΩ and 6–9 MΩ, respectively. Pipettes were backfilled with intracellular solution (in mM, 130 K-gluconate, 0.1 EGTA, 10 HEPES, 7 NaCl, 0.3 MgCl_2_, 2 ATP, 0.5 GTP, 5 phosphocreatine; 280 mOsm). Intracellular solutions were balanced to a pH of 7.3 with 10 M KOH and osmolality was cross-checked to fall within the range of 275–290 mOsm. Data were acquired at 10 kHz using Clampex software (PClamp 10.4; Molecular Devices).

We examined intrinsic properties of motoneurons and ventral interneurons and terminated experiments if access resistance was greater than 25 MΩ for motoneurons and 35 MΩ for interneurons, if cells had a resting membrane potential greater than −50 mV, or if action potential amplitude was less than 60 mV at baseline. Cells were excluded from analysis if access resistance deviated by more than 20% by the end of the recording. We held all cells at a membrane potential of approximately −75 mV during experiments, after correcting for a liquid junction potential of 14.3 mV. The liquid junction potential was calculated in Clampfit (PClamp 10.4; Molecular Devices), using the ionic composition of our extracellular and intracellular solutions. Protocols for the examination of intrinsic properties have been described elsewhere [82].

### Pharmacology

Dopamine hydrochloride (Sigma-Aldrich, Inc., St. Louis, MO) was bath applied in separate experiments at 1 µM, 3 µM, 10 µM, 30 µM, 100 µM, and 300 µM to determine dose-dependent effects on motor activity. The receptor-selective agonists we used included SKF 81297 for D_1_-like receptors (10–50 µM; Tocris, Minneapolis, MN); quinpirole for D_2_-like receptors (10–50 µM; Tocris); and the D_1_/D_2_ receptor co-agonist SKF 83959 (10–50 µM; Tocris). For dopamine receptor antagonists we used the D_1_-like antagonist SCH-23390 (10 µM; Tocris); the D_2_-like antagonists sulpiride (20 µM) and L-741,626 (12 µM); the selective D_3_ receptor antagonist SB 27701A (5 µM; Tocris); the selective D_4_ receptor antagonist L-745, 870 (5 µM; Tocris). We also used the α_2_ adrenergic receptor antagonist, yohimbine (2–4 µM; Tocris). Endogenous dopamine levels were manipulated with the DAT inhibitor GBR-12909 (10 µM; Hello Bio, Princeton, NJ) and the monoamine oxidase A & B inhibitor bifemelane (50 µM; Tocris).

### Immunoprecipitation for D_1_ and D_2_ receptors

Spinal cords were dissected in ice-cold (4–8°C) aCSF and homogenized in lysis buffer containing 50 mM TrisHCl, 150 mM NaCl, 10 mM EDTA, 0.1% Triton-X, and 5% Glycerol. Lysis buffer contained protease inhibitors (Sigma) and phosphatase inhibitors (GBiosciences). We homogenized three spinal cords in 100 µL of buffer and incubated them on ice for 1 hour before centrifuging them at 10,000 rpm for 30 minutes at 4°C. Lysates were then extracted and stored at −20 °C.

To reduce nonspecific binding, we first incubated lysates in anti-rabbit Ig agarose beads (Trueblot; Rockland Inc., Limerick, PA) for 30 minutes, on ice and in the absence of primary antibody. We then removed the supernatant and incubated the lysates on ice for 1 hour with rabbit antibody to D_2_ receptors (1 μg per 100 μL, Millipore). Anti-rabbit Ig IP beads were added and samples were incubated overnight at 4°C with gentle agitation. Immunoprecipitates were washed with lysis buffer, heated in loading buffer (350 mM Tris, 30% glycerol, 1.6% SDS, 1.2% bromophenol blue, 6% β-mercaptoethanol) to 95°C for 10 min, electrophoresed on a precast SDS gel (4–12% Tris HCl; BioRad, Hercules, CA), and transferred onto a nitrocellulose membrane. After blocking, the membranes were incubated with guinea pig antibody to D_1_ receptors (1:400; Alomone, Jerusalem, Israel) and rabbit antibody to D_2_ receptors (1:500; Millipore, Burlington, MA), washed, incubated for 2 hours at room temperature in fluorophore-conjugated secondary antibodies (IRDye anti-guinea pig and Trueblot anti-rabbit IgG DyLight, 1:1000), and visualized via antibody fluorescence at 680 or 800 nm (Odyssey CLx; LI-COR Biosciences, Lincoln, NE).

### Post hoc verification of motoneurons

Following the completion of experimental protocols, we post-fixed slices overnight in 4% paraformaldehyde (PFA) at 4°C, washed them the next morning in phosphate-buffered saline (PBS) for three 20-minute intervals, and stored them at 4°C. On the day we performed immunohistochemistry, we first washed the slices for 30 minutes (3 × 10 minutes) in PBS with 0.5% Triton-X 100 (PBST) then incubated them at room temperature (21-23°C) for 6 hours in blocking solution containing 10% donkey serum in 0.5% PBST. Primary antibodies for choline acetyltransferase (ChAT) (1:500; goat anti-ChAT; Millipore Cat No. AB144) and the fluorescence marker (1:1000; rabbit anti-FITC, ThermoFisher Cat No. 11090) were diluted in blocking solution and the slices incubated with them for at least 24 hours at room temperature. Slices were then washed in PBST (0.5%) for 80 minutes (4 × 20 min) at room temperature before secondary antibodies (donkey anti-rabbit Alexa Fluor 1:1000, Life Technologies Cat No. A-21206; Donkey anti-Goat Alexa 568 1:1000; Cat No. A21447, Life Technologies, Carlsbad, CA) were applied for 4–8 hours at room temperature, then washed for 80 minutes in PBST (0.5%). Slices were mounted and coverslipped with fluorescent mounting medium (Vectashield; Vector Laboratories, Burlingame, CA); coverslips were separated from slides by 500 μm spacers to prevent crushing.

### Imaging

All sections processed via immunohistochemistry were imaged on a Nikon A1R MP_+_ microscope (Nikon, Tokyo, Japan) operating in confocal mode with a 16× water-immersion objective lens (numerical aperture [NA] = 0.8, working distance [WD] = 3 mm). Image acquisition used a z-step of 1 µm and averaged two frames with a resolution of 2048 × 2048. Pixel dwell time was 2.5 ms and exposure settings were maintained for all sections. We used NIS-Elements software (Nikon) for image acquisition and ImageJ to perform maximum intensity projections of z-stacks.

### High-performance liquid chromatography

Monoamine content of neonatal and adult spinal cords was measured using high-performance liquid chromatography (HPLC). We dissected spinal cords from neonatal (P3, n = 11) C57BL/6 mice in aCSF as described above and extracted adult spinal cords (P60, n = 17) with a pressure ejection method. Tissue was then flash-frozen with liquid nitrogen, stored at −80°C, and analyzed for biogenic amines with a modified version of the Parent et al. HPLC method [76]. Tissue was homogenized in ice-cold 0.1 M perchloric acid. We centrifuged the homogenate and used 10 μl of supernatant in the assay, employing an Atlantis dC18 column (Waters, Milford, MA) and an electrochemical detector.

### Data analysis

We determined the relative inhibitory or excitatory effects of dopamine and dopamine receptor agonists on spontaneous motor network activity using methods similar to those in our previous work [23]: we calculated a response ratio from single ventral root neurograms between the root mean square of 5 minutes of basal spontaneous activity and 5 minutes of activity recorded 20 minutes after adding the drug. We subtracted 1 from the response ratio so that positive values reflect excitation and negative values reflect inhibition. The response ratio was used as a high throughput assay to detect global changes in network activity. Neurogram data were analyzed with Spike2 software. Bursts of spontaneous activity were analyzed using Clampfit (Molecular Devices) to determine how episode number and amplitude contributed to changes in response ratio. Spectral analyses were conducted using Spinalcore software [83] whenever we detected excitatory changes or observed rhythmicity.

Patterns of rhythmic motor activity recorded from single ventral roots were analyzed with autowavelet spectral analysis. We created and analyzed frequency–power spectrograms by selecting regions of interest around frequency ranges that coincided with those in the raw neurogram. The spectrograms revealed two high power regions that reflect distinct rhythms evoked by dopamine at high concentrations: a slow 0.01–0.04 Hz rhythm and a fast 0.8–1.2 Hz rhythm. Regions of interest were selected within these frequency ranges and analyzed over the time course of each experiment. Frequency power within selected regions of interest that corresponded to the fast and slow rhythms were used as a measure of rhythm robustness. Data were segmented into 30 s bins and averaged over 5-minute intervals for statistical analysis. We used tools available in Spinalcore for all analyses of rhythmic motor activity [83], consistent with Sharples and Whelan [13].

Motoneuron and interneuron intrinsic properties measured during whole-cell patch-clamp experiments were analyzed as in our previous work [82,84], with the exception of repetitive firing analyses, which examined the instantaneous firing rate for the first spike interval and steady-state firing separately.

### Experimental design and statistical analysis

All experiments were repeated measures. We tested for differences in the magnitude of effects between conditions with one-way ANOVAs, focusing on comparisons to time-matched vehicle controls. Two-way ANOVAs compared baseline to multiple post-drug conditions. All effects surpassing a significance threshold of p < 0.05 were further examined with post hoc analyses. We used Holm-Sidak post hoc tests to compare all treatment conditions to the appropriate normalized time-matched vehicle control. Data that violated assumptions of normality (Shapiro–Wilk test) or equal variance (Brown–Forsythe test) were analysed via nonparametric Mann–Whitney *U* (if two groups) or Kruskal–Wallis (if more than two groups) tests.

## Supplemental Material

**Supplemental Figure 1:**
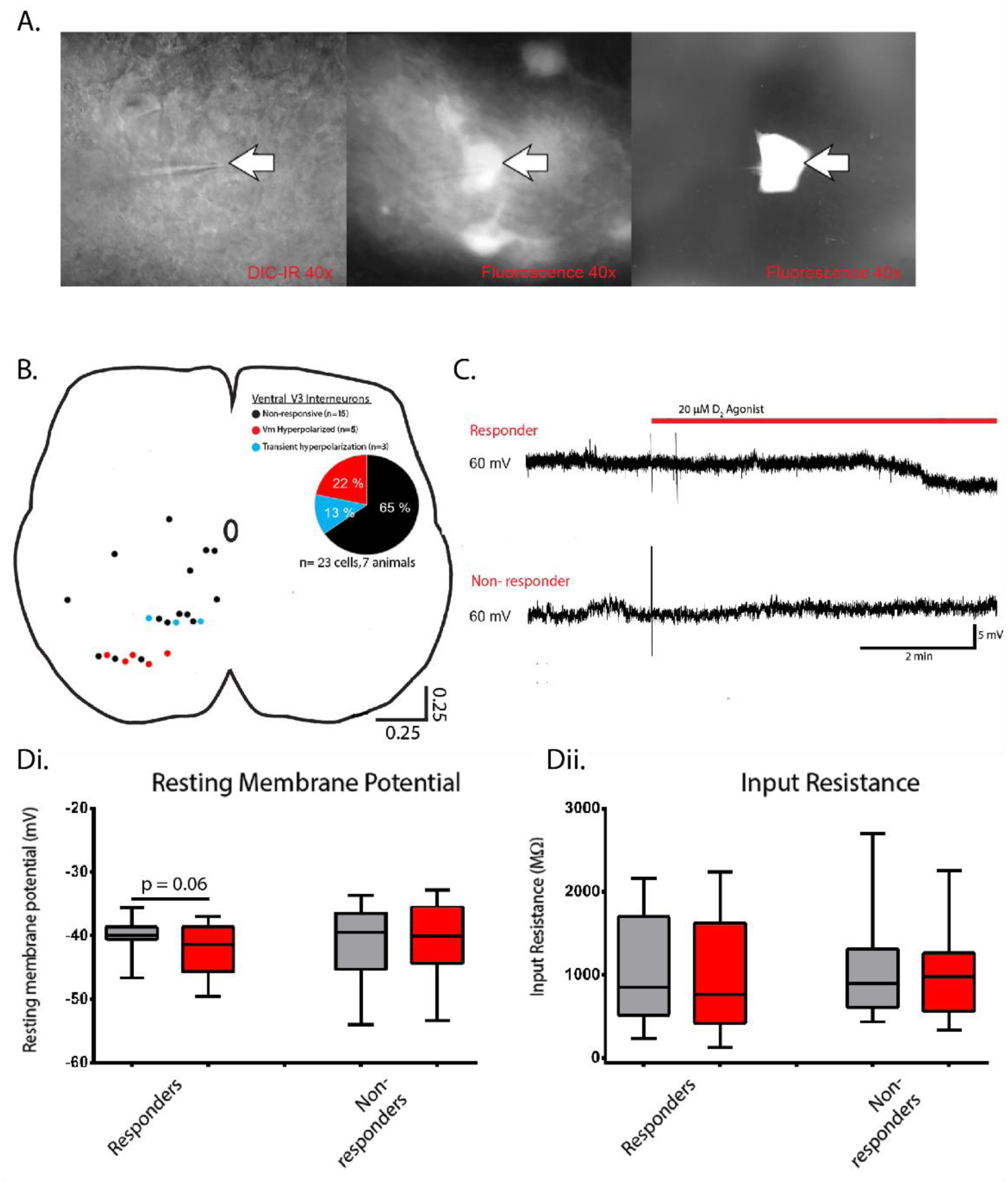
D_2_-receptor control of V3 interneurons. Whole-cell patch-clamp recordings obtained from lamina V3 interneurons. A. Visually guided patch showing pipette tip on a GFP labelled-V3 interneuron. B. V3 interneuron location was measured relative to the central canal and X-Y positions normalized to the distance of ventral and lateral borders of lumbar slices. Non-responders represented by black dots, responders by red dots, and transient responders by blue dots. C. A sample trace showing a hyperpolarization of the resting membrane potential following administration of 20 µM quinpirole in responders (top trace) and non-responders (bottom trace). Di. Box and whisker plots showing a change in resting membrane potential in responders and non-responders before (gray) and after (red) administration of quinpirole. Dii. Input resistance graphed in a similar manner as Di.

**Supplemental Video 1:** Spontaneous movements in neonatal (P3) C57 mice.

